# EMU: reconfigurable graphical user interfaces for Micro-Manager

**DOI:** 10.1101/2020.03.18.997494

**Authors:** Joran Deschamps, Jonas Ries

## Abstract

Advanced light microscopy methods are becoming increasingly popular in biological research. Their ease of use depends, besides experimental aspects, on intuitive user interfaces. The open-source software Micro-Manager offers a universal interface for microscope control but requires implementing plugins to further tailor it to specific systems. Since even similar devices can have different Micro-Manager properties (such as power percentage versus absolute power), transferring user interfaces to other systems is usually very restricted.

We developed Easier Micro-Manager User interface (EMU), a Micro-Manager plugin, to simplify building flexible and reconfigurable user interfaces. EMU offers a choice of interfaces that are rapidly ready to use thanks to an intuitive configuration menu. In particular, the configuration menu allows mapping device properties to the various functions of the interface in a few clicks. Exchanging or adding new devices to the microscope no longer requires rewriting code. The EMU framework also simplifies implementing a new interface by providing the configuration and device interaction mechanisms. The user interface can be built by using a drag-and-drop tool in one’s favorite Java development environment and writing a few lines of code for compatibility with EMU.

Micro-Manager users now have a powerful tool to improve the user experience on their instruments. EMU interfaces can be easily transferred to new microscopes and shared with other research groups. In the future, newly developed interfaces will be added to EMU to benefit the whole community.

## Background

Light microscopy is an ever-growing field with countless applications in biosciences. A substantial portion of technology developments and cutting-edge research are carried out on custom microscopes because of their high flexibility. Beyond the mechanical and optical requirements of such microscopes, researchers face the challenges of controlling the hardware and presenting the users with an intuitive interface. LabView (National Instrument) has historically been one of the main go-to solutions thanks to its rapid prototyping capabilities and its compatibility with a wide range of commercial hardware devices. However, LabView is an expensive proprietary software and is difficult to maintain or share. The code is then often being tailored to one application in particular. A popular alternative to LabView in the light microscopy field is Micro-Manager (μManager) (1), an open-source software based on ImageJ (2).

μManager is a ready-to-use platform compatible with many popular hardware devices. The support for new devices is community driven and constantly expanding. In addition, μManager features the possibility to run scripts or plugins to perform tailored experiments. It offers a universal interface allowing image acquisition and interaction with the microscope devices. In μManager, each device is controlled through its device properties, for instance laser power percentage or filter wheel position. While some device properties are in principle intuitive, others can be too specialized or irrelevant to the user. Therefore, several tools are available to simplify the microscope control for non-specialists, such as bundling properties together, hiding their values, and presenting only self-explanatory names to the user (configuration preset groups tool) or a window that can be customized with action-triggering buttons (quick-access panel). These tools are easy to set-up and use; however, because they are aimed at covering general needs, they cannot rival with a tailored interface in terms of user experience.

The preferred way to implement a custom interface in μManager is by writing a plugin, which is automatically detected by μManager at start-up. The plugin can then be loaded using the main μManager window. Several dedicated libraries exist in Java in order to build graphical interfaces, for instance the widely used Swing (3). Working with the Swing toolkit requires experience and knowledge of the library. Fortunately, most major Java integrated development environments (e.g.: Eclipse, Netbeans, IntelliJ) provide graphical tools to build a user interface (UI). With a basic understanding of Swing, users can assemble complex UIs by placing (“drag-and-drop”) components on a panel or a frame without writing a single line of code. Nonetheless, programming is then required to call μManager methods and modify device states when the user interacts with the UI components.

Yet, similar devices might have slightly different properties (e.g. laser power percentage vs absolute laser power) or different property values (e.g. “On” vs “1”) depending on the manufacturer or the device adapter implementation. Therefore, care should be taken to make interfaces reconfigurable and flexible enough to accommodate new devices without rewriting code. Otherwise, the interface will not be transferable to similar systems or other research groups, and expert knowledge will be required whenever a device changes on the microscope. Regrettably, interfaces are usually bound to a single system and never benefit other users.

We developed Easier Micro-Manager User interfaces (EMU) to simplify implementing reconfigurable and easily transferable UIs for μManager, as well as to offer a repository of already existing interfaces for the community to use.

## Results and Discussion

### Implementation

EMU is a μManager (2.0.0-gamma) plugin, written in Java (1.8), that includes its own plugin system. It allows users to load an EMU plugin from a list of available interfaces, ether one of the examples or a custom-made UI. In particular, users have access to an intuitive menu to configure the UI. The configuration menu displays a list of parameters that can be modified to tailor the plugin to the microscope.

#### EMU main classes

The EMU framework is based on the Java Swing toolkit (3). Its own plugins (i.e. user interfaces) consist of multiple panels (*ConfigurablePanel* class) arranged within a single frame (*ConfigurableMainFrame* class), as illustrated in figure 1. These two classes are extensions of the Swing classes *JPanel* and *JFrame* respectively.

**Fig. 1.**
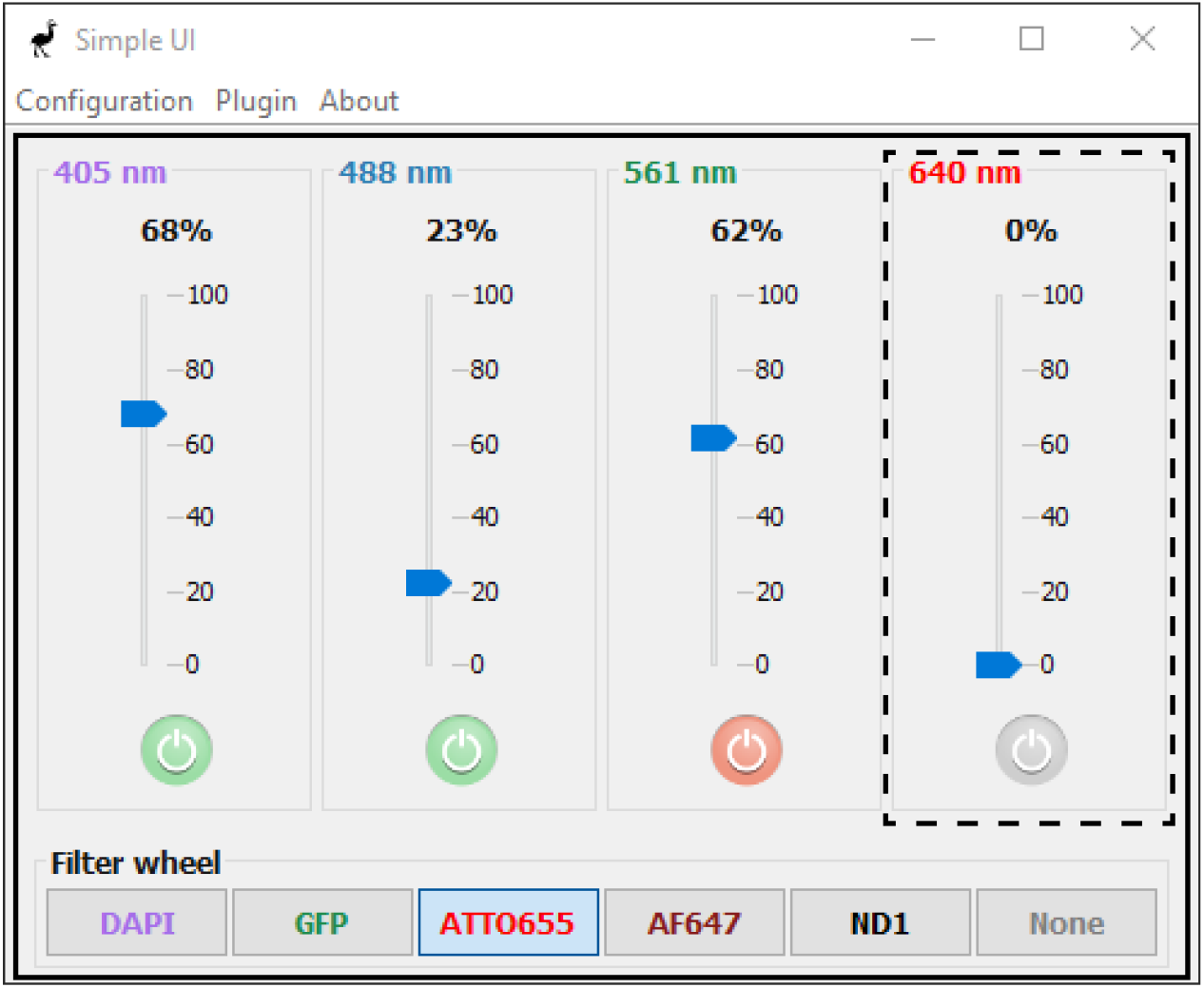
EMU plugin example. SimpleUI, one of the interfaces available by default within EMU, can control four lasers and an optional filterwheel. The solid box delimits the *ConfigurableMainFrame* instance while the dashed box is one of the *ConfigurablePanel* objects of the interface. All titles and colors are UI parameters, while the UI properties are the laser power percentage and on/off (for each laser) and the filter wheel position. The on/off buttons can be disabled. The interface has a single plugin setting: showing or hiding the filter wheel panel.

*ConfigurablePanel* objects are units of device control and can declare UI properties (*UIProperty* class) and UI parameters (*UIParameter* class). UI properties are aimed at being paired with a μManager device property. Several types of UI properties are available in EMU (see table 1), among which properties with a fixed number of states (e.g. on/off) or with the possibility to rescale the device property value (e.g. use a laser power percentage in the UI to modify an absolute power device property). User interactions with the graphical components (e.g. buttons) trigger UI properties, which in turn change the state of the corresponding device properties and therefore of the hardware devices themselves. Using UI properties with constraints simplifies their interplay with the graphical components. For instance, a toggle button (selected/unselected) is well suited to trigger a UI property with only two states (*TwoStateUIProperty* class). UI parameters, on the other hand, change the look and layout of the *ConfigurablePanel* objects (e.g. text or color). Table 2 summarizes the UI parameters provided by the EMU framework. All *ConfigurablePanel* objects are instantiated in the plugin’s *ConfigurableMainFrame* instance. The latter also declares plugin settings (*Setting* class), which can be for instance an option to hide or show a *ConfigurablePanel* (see table 3 for a list of setting types).

**Table 1.** *UIProperty* class and child classes. A *UIProperty* object only changes the state of a device property within the property limits or allowed values. *UIProperty* child classes have additional constraints, as specified in the second c olumn. In the EMU configuration me nu, these constraints lead to additional fieds as indicated in the third column.

| Class | Note | in EMU configuration menu |
| --- | --- | --- |
| *UIProperty* | General UI property | device and property fields |
| *SingleStateUIProperty* | Accepts a single-state | + field for the state value |
| *TwoStateUIProperty* | Accepts an *On* and an *Off* state | + fields for the *On* and *Off* values |
| *MultiStateUIProperty* | Accepts a fixed number of states | + field for each state value |
| *RescaledUIProperty* | Rescales value to $slope \cdot v + offset$ | + fields for the *slope*/*offset* values |

**Table 2.**
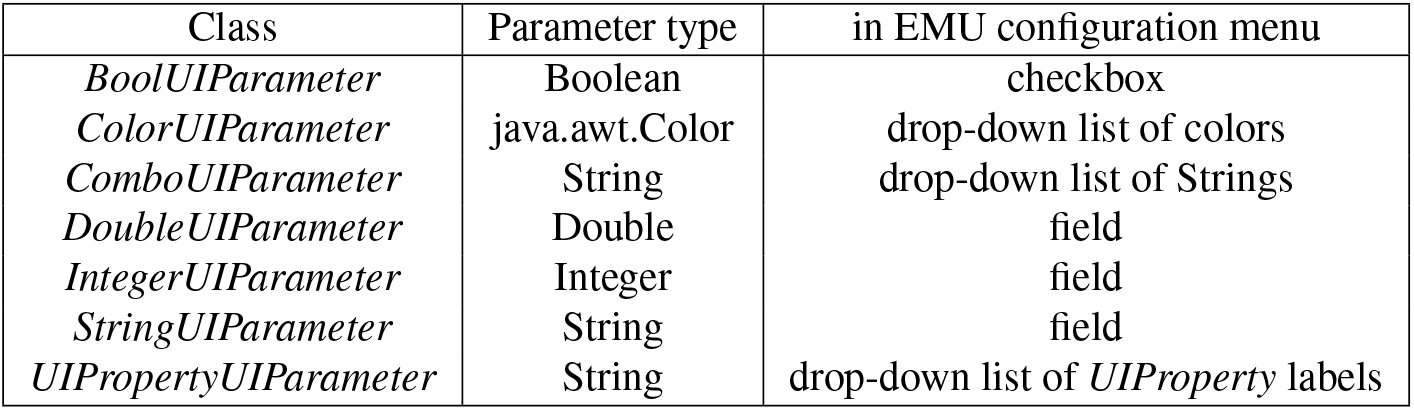
*UIParameter* child classes. Each *UIParameter* child class holds a member variable represented by the type indicated in the second column. In the EMU configuration menu, the UI parameters appear as specified in the third column.

**Table 3.**
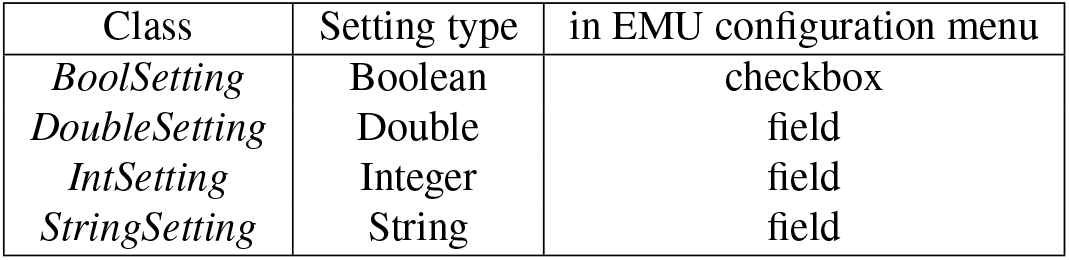
*Setting* child classes. Each class holds a member variable represented by the type indicated in the second column. All *Setting* child classes appear in the EMU configuration menu as a field, except for the *BoolSetting* settings, which appear as checkboxes.

| Class | Setting type | in EMU configuration menu |
| --- | --- | --- |
| *BoolSetting* | Boolean | checkbox |
| *DoubleSetting* | Double | field |
| *IntSetting* | Integer | field |
| *StringSetting* | String | field |

Upon starting EMU, the plugin settings, UI properties and UI parameters are automatically aggregated and given a value based on the last known configuration. They appear in the configuration menu and their values can easily be changed each time a device is exchanged or added to the microscope.

#### Implementing an EMU plugin

In order to build a user interface compatible with EMU, programmers must create their own extensions of the *ConfigurablePanel* and *ConfigurableMainFrame* classes. In particular, this requires laying out graphical components using the Swing toolbox to assemble the desired interface. Note that this can be achieved using drag-and-drop tools offered by most modern development environments.

Both *ConfigurableMainFrame* and *ConfigurablePanel* classes inherit few methods that must be implemented. These methods include the declaration of the settings (plugin setting, UI properties and UI parameters), as well as methods called when UI properties/parameters have changed so as to reflect these changes in the relevant graphical components state. Finally, the effect of user interactions with the components should be implemented using action/state listeners from the Swing library. To that purpose, EMU provides static methods in the *SwingUIListeners* class, covering usual cases and thus simplifying the implementation of *ConfigurablePanel* child classes.

EMU uses the Java *ServiceLoader* mechanism to import its plugins. Therefore, EMU plugins should also include an implementation of the *UIPlugin* class and be compiled in a .jar file, following the guidelines of the *ServiceLoader* framework.

#### Loading a plugin into EMU

Compiled .jar files should be placed in the μManager installation folder under the “EMU” folder. At start, EMU automatically detects the plugin and adds it to its list of available user interfaces.

### Interface

EMU can be started in μManager from the plugin menu. Upon the first start, users can choose a UI among the detected interfaces. They are subsequently presented with a configuration menu. In the latter, four tabs give access to the different configurable settings: plugin settings, UI properties, UI parameters and global settings. Plugin settings and UI parameters alter the look and default values of components in order to tailor the interface to the system and make it more intuitive to users. This ranges from hiding irrelevant panels to giving self-explanatory titles to subpanels. Figure 2 shows the parameters configuration menu in the case of SimpleUI (figure 1), one of the example UIs in EMU. Global settings are EMU options, such as displaying warning messages when misconfigured.

**Fig. 2.**
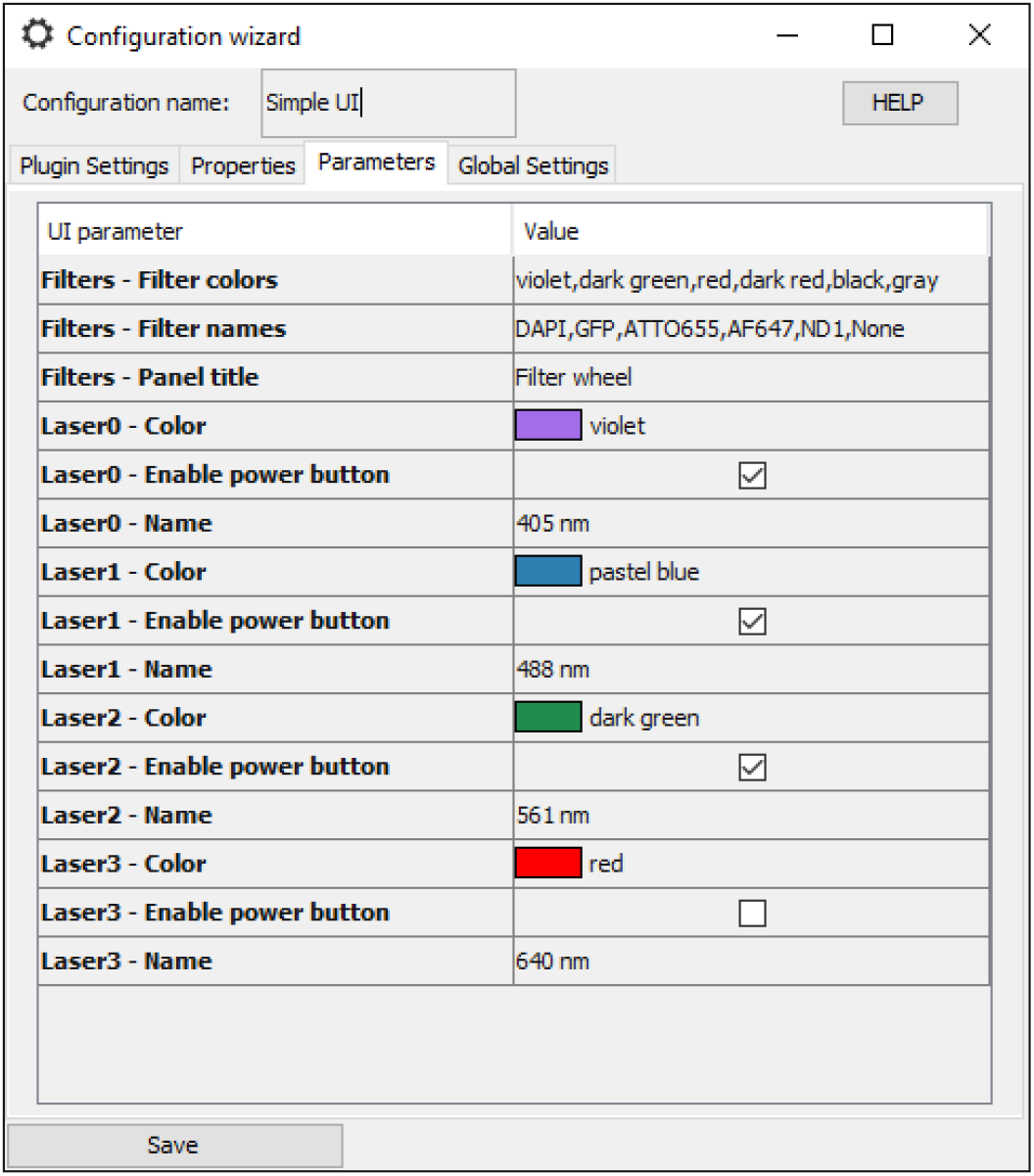
Configuring the UI parameters. In the configuration wizard, users can modify the interface UI parameters. Here shown for SimpleUI.

The most important configurable aspects are the properties. In the configuration menu, users can map the UI properties to μManager device properties by selecting first a device, then the pertinent device property (see figure 3 in the case of SimpleUI). As some UI properties have states (see table 1), users must also enter suitable device property values. The latter are easily found in the device property browser of μManager. Thereafter, whenever users interact with the graphical components (e.g.: button), the corresponding UI property will change the state of the device property accordingly.

**Fig. 3.**
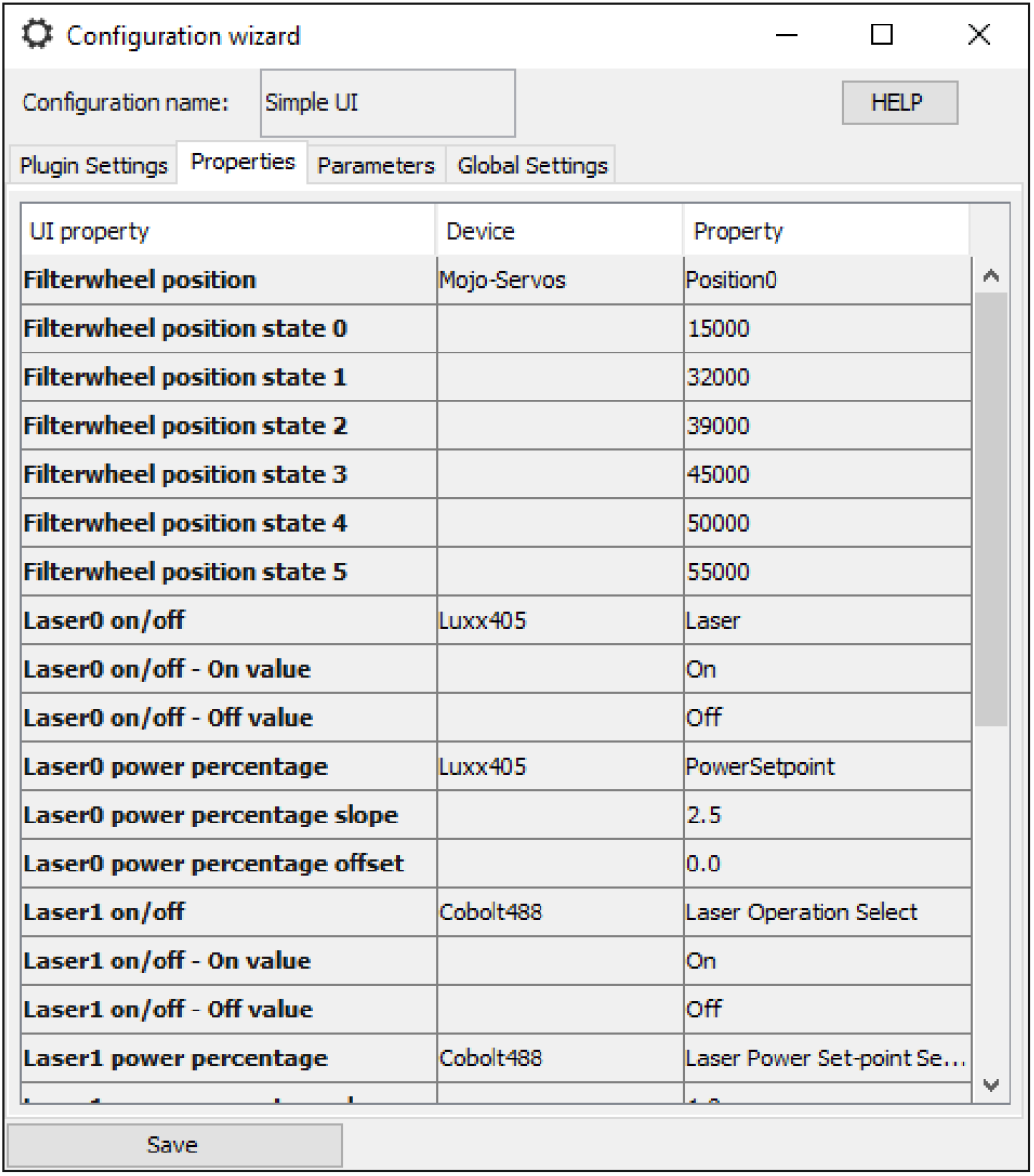
Configuring the UI properties. In the properties tab, users can map devices and the relevant properties to the interface UI properties. Here shown for SimpleUI.

After saving the configuration, a human-readable file is automatically created in the μManager installation folder (under/EMU/). Multiple configurations can coexist and be saved in the configuration file. The EMU menu (see the menu bar in 1) allows users to switch between UIs or between configurations in a single click. Finally, at each start of EMU, the configuration file is loaded and the last known plugin configured.

### Example cases

To illustrate the flexibility of EMU plugins, we can consider a simple example such as the UI shown in figure 1: SimpleUI. The plugin controls four lasers, allowing users to turn their emission on and off and change their power percentage. Laser names and colors can be set in the configuration menu to describe their intended use (see figure 2). Several cases can be encountered when working with lasers in μManager: (i) all lasers have a laser emission and a power percentage device property, (ii) some lasers have an absolute power instead of a power percentage device property and (iii) some lasers do not have a laser emission device property. In (i), the UI properties can be mapped to the device and relevant device properties. Since the UI properties in charge of changing the laser powers are *RescaledUIProperty* objects (see table 1), the user can leave the *slope* and *offset* values (see figure 3) equal to 1 and 0 respectively. For the lasers in (ii), the laser percentage UI property is mapped to an absolute power device property. There, the *slope* parameter should be set to *max/*100 (with *max* being the maximum laser power), effectively rescaling the interface power percentage to the range {0*, max*} of the device property. Finally, for lasers in the case (iii) (such as the laser diodes described in (4)), no laser emission device property exists. The laser emission UI properties should remain unconfigured. Using the UI parameters, the on/off buttons can be disabled in the interface. In all three cases, the interface can be configured to reflect an accurate picture of the microscope lasers.

The example plugins currently available in EMU are simple. However there is no limit on how complex an EMU plugin can be. We routinely use htSMLM (5) (see Additional file 1), an EMU plugin aimed at controlling a wide-field microscope. htSMLM features four lasers, two optional iBeamSmart lasers (Toptica), an axial focusing panel, up to four filter wheels and multiple toggle buttons. It also includes tools for localization microscopy (6–8) (automated activation script) and for performing different type of acquisitions in series (localization microscopy, multi-slice localization, snapshot, time series, z-stacks) that take into account the device properties linked to the interface. This interface has been used by scientist with a wide variety of backgrounds and projects, ranging from biology to optics (9–12).

## Discussion

Well-designed user interfaces are key components of microscopes. Not only do they facilitate the use of the instrument, but they accelerate research by ensuring an efficient interaction between the users and the hardware. While μManager offers a universal interface and some customization tools, complex instruments require tailored interfaces. Several user interfaces are available to control microscopes with μM, such as a Matlab-based UI (13) for the Olympus IX83, the interfaces for the OpenSPIM (14), OpenSpinMicroscopy (15) and diSPIM (16) (ASIdiSPIM plugin in μManager) microscopes. Because these interfaces perform specialised acquisitions and hardware control, they are meant to control very specific hardware devices. Henceforth, any departure from the original devices requires modifying the source-code. In contrast, the references to the hardware devices are not hard-coded in EMU plugins. Indeed, they are supplied by the users when configuring the interface, along with the property values. A direct consequence is that similar microscopes can be controlled by the same interface, as long as their hardware devices are compatible with μManager.

Configuring an EMU plugin to work with a microscope only takes a couple of minutes. In the same way, exchanging devices on the set-up is equally convenient, as users only need to start the EMU configuration menu and change the relevant lines to reflect the presence of the new device or new state. Because no device property or device property value is hard-coded, easy sharing and transfer of user interfaces is streamlined between similar microscopes as well as between research groups.

While EMU simplifies building specialised user interfaces and is compatible with drag-and-drop tools, programming knowledge is nonetheless required to implement the mandatory EMU methods. In order to reduce the difficulty of the task, we provide online the EMU guide (17), which includes a user and a programming guide, a step-by-step tutorial on how to create an EMU plugin, as well as multiple examples of the use of EMU classes.

Since EMU was developed with the aim of creating easily shareable interfaces, we intend to integrate a number of new plugins to the source-code as they become available. Currently, EMU includes only SimpleUI and a plugin controlling iBeamSmart lasers from Toptica. In the future, htSMLM will be integrated as well, along with any contribution from other users.

## Conclusions

μManager is a successful open-source software and is widely used to control custom microscopes. However, its universal interface does not provide a user experience comparable to tailored UIs and many research groups lack the knowledge to implement such interfaces. EMU provides ready-to-use interfaces and is now distributed with μManager 2.0.0-gamma. Using the EMU framework also simplifies writing UIs for μManager as it includes an advanced configuration system and takes care of all interactions with device properties. Implementing an EMU plugin can be done using graphical tools available in most development environments, with only a few methods to implement oneself in order to obtain a functional interface. Thereafter, users have access to an intuitive configuration menu to set-up the UI for their system. We hope that in the future users will contribute their own UI to EMU in order to benefit the whole community.

## Abbreviations

EMU: Easier Micro-Manager User interfaces
μManager: Micro-Manager
UI: user interface

## Acknowledgements

We would like to thank Serge Dmitrieff, Philipp Hoess, Robin Diekmann and Ingmar Schoen for comments on the manuscript.

## Author’s contributions

J.D. and J.R. designed the project, J.D. implemented and tested the software.

## Additional Files

**Additional Figure 1.**
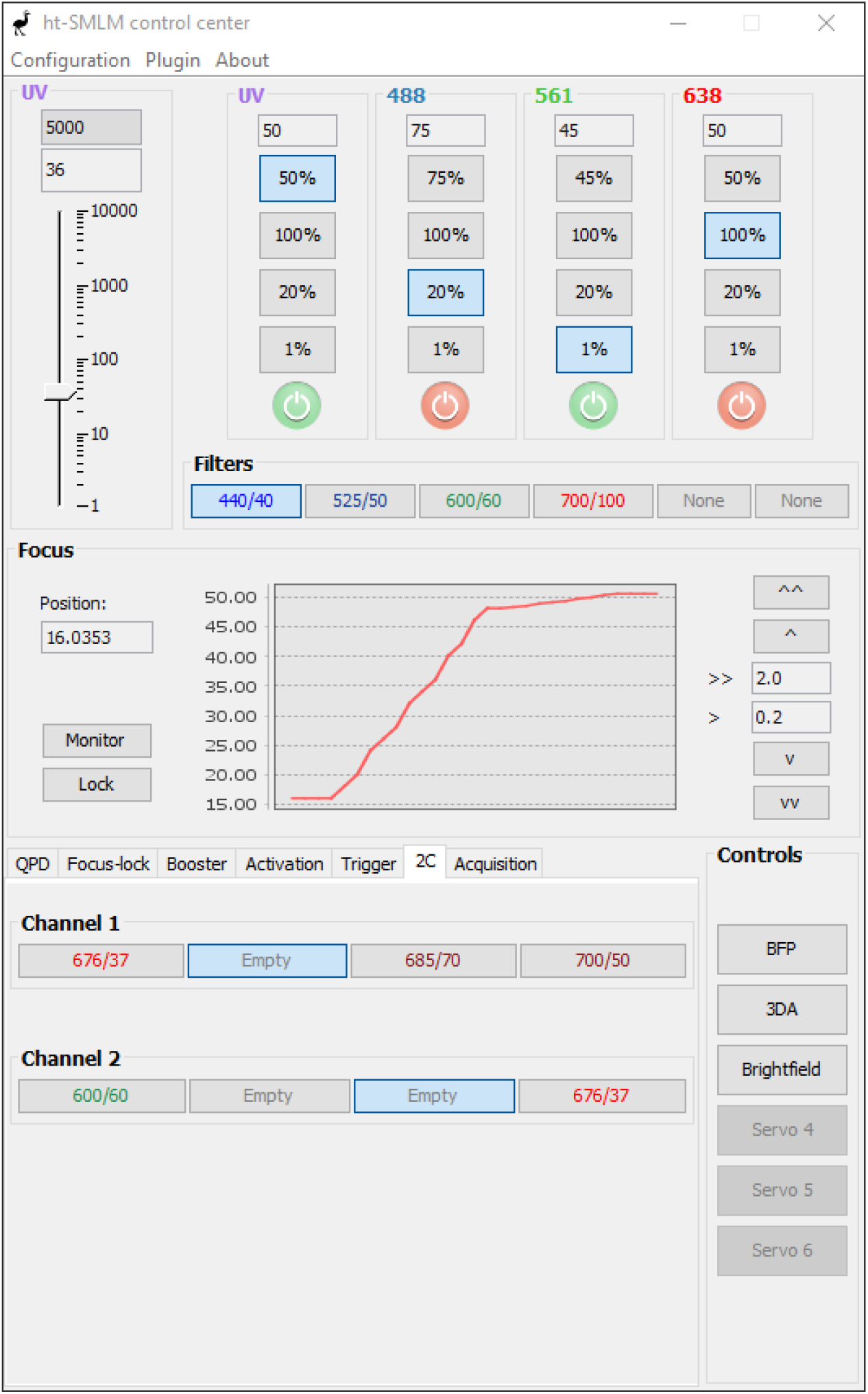
htSMLM interface. htSMLM is a complex EMU plugin aimed at controlling a wide-field or localization microscope. Besides the controls for multiple lasers, filter wheels and focus, it features tools to perform series of experiments sequentially (e.g. localization microscopy, time series, z-stack…etc…).

